# EFFECT OF LAND TENURE ON ADOPTION OF CLIMATE CHANGE ADAPTATION. EVIDENCE FROM MALAWI

**DOI:** 10.1101/2020.12.03.409615

**Authors:** Horace Phiri

## Abstract

The definitive aim of this study was to bring to fore the evidence of the importance of tenure considerations in the designing, development, and implementation of climate change programs. This was done by analyzing how land tenure affects the use of adaptation strategies in Malawi. Using secondary data from the Integrated Household Survey (IHS4), a multinomial logit model was fitted to analyze determinants of adoption of climate change adaptation strategies. Land tenure has shown to significantly affect the adoption of the technologies in question. Insecure arrangements such as borrowing and renting land tend to discourage adoption. The proliferation of borrowed or rented in Malawi’s agricultural sector necessitates intervention to encourage adaptation on those farms to avoid land degradation.

## 1 Introduction

Globally, the relationship between farm level and land tenure security is one of the widely studied research questions (Leonhardt, Penker, & Slahofer, 2019). Theoretically, secure property rights raise farmers’ propensity to invest (Deininger, 2003) the lack of which as is the case in customary land tenure systems hampers long term investment (Place & Otsuka, 2001). Despite these assertions, empirical evidence has been non-conclusive. Investment in soil conversation exhibited a positive relationship with land tenure security (Ali, Deininger, & Goldstein, 2011; Gebremedhin & Swinton, 2003; Lovo, 2016). Contrary to common sense and earlier empirical work results obtained by Brasselle, Gaspart, and Platteau (2002) cast doubt on the existence of a systematic influence of land tenure security on investment. Similarly, an earlier study conducted by Place and Hazell (1993) in Ghana, Kenya, and Rwanda concluded that land rights are not a significant factor in determining investments in land improvements.

Overall, the literature indicates that land tenure has the potential to impact farm incomes through its impact on farm-level investment in technologies (Lawry et al., 2017). The enacting of the amended Land Act (2016) in Malawi and the increasing usage of rented land raises fresh concerns about how the changing land access arrangements will impact farm investment. Therefore, the definitive aim of this study was to examine the impact of tenure on the choice of climate change adaptation. This was done to bring forth evidence of the importance of tenure considerations in the designing, development, and implementation of climate change programs in Malawi.

## 2 Econometric Framework and Data sources

### 2.1 The analytical framework of the study

Farm household choice of whether or not to use any climate change adaptation option could fall under the general framework of utility and profit maximization. Consider a rational farmer who seeks to maximize the present value of expected benefits over a specified time horizon, and much choose among a set of *J* adaptation options. The farmer *i* decides to use *J* adaptation option if the perceived from option *J* is greater than the utility from other options (say, *k*) depicted as

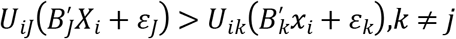

Where *U*_*iJ*_ *and U*_*ik*_ are the perceived utility by farmer *i* of the adaptation options *j* and *k*, respectively; *X*_*i*_ is a vector of explanatory variables that influence the choice of adaptation option; *B*_*j*_ and *B*_*k*_ are parameters to be estimated; and *ɛ_J_* and *ɛ_k_* are error terms. Under the revealed preference assumption that the farmer practices an adaptation option that generates net benefits, and does not practice an adaptation option otherwise, we can relate the observable discrete choice of the practice of the unobservable (latent) continuous net benefit variable as *Y*_*ij*_ = 1 if *U*_*iJ*_ > 0 and *Y*_*ij*_ = 0 if *U*_*iJ*_ < 0. In this formulation, *Y*is a dichotomous dependent variable taking the value 1 when the farmer chooses an adaptation option in question and 0 otherwise. Accordingly, the probability that the farmer i will choose adaptation option j among the set of adaptation options could be defined as follows.

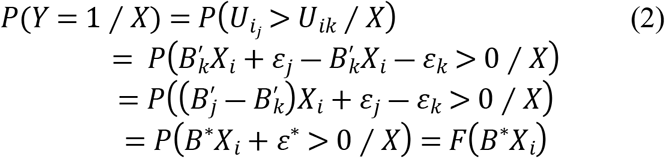

Where *ɛ*\* is a random disturbance term, *B*\*is a vector of unknown parameters that can be interpreted as the net influence of the vector of explanatory variables influencing adaptation, and *F* (*B*\**X*_*i*_) is the cumulative distribution of *ɛ*\* evaluated at *B*\**X*_*i*_.

In this study we had four adaptation choices, the multinomial logit (MNL) model was used to analyze the behavior of farmers. Thus the probability that household i with characteristics X chooses adaptation option j specified as follows;

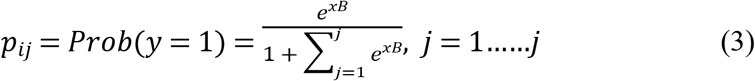

Where *B* is a vector of parameters that satisfy *ln*(*P*_*ij*_ / *P*_*ik*_) = *X*′(*B*_*j*_ − *B*_*k*_) (Greene, 2003). Unbiased and consistent parameter estimates of the MNL model in equation (3) require the assumption of independence of irrelevant alternatives (IIA) to hold. The IIA assumption requires that the likelihood of a household using a certain adaptation measure needs to be independent of the alternative adaptation measures used by the same households. Hausman Specification Test is used to ascertain the validity of the IIA assumption(Hausman, 1978).

Data used in the study was from the Fourth Integrated Household Survey 2016/17 (IHS4) conducted by the Malawi National Statistical Office in collaboration with the World Bank Living Standards Measurement Survey Team. The IHS4 is the fourth cross-sectional survey in the IHS series and was fielded from April 2016 to April 2017. The IHS4 2016/17 collected information from a sample of 12,447 households, representative at the national-, urban/rural-, regional- and district-levels.

### 2.2 Variables and descriptive statistics

### 2.3 Dependent variables

Our dependent variable is the adoption of climate change adaptation technologies. A wide range of innovations is available for use by farm families to cushion the impact of climate change. Based on the known impacts of climate change in this study we consider three broad categories of technologies: Erratic Rainfall (ER). Soil Fertility (SF) and A combination of the two groups (SFER). We only considered the technologies already adopted and not the intention to adopt. Almost two-thirds of participants reported using SFER, 21.5% had SF managing practices on their fields and 11% had innovations to minimize the impact of ER.

### 2.4 Independent variables

Descriptive statistics for independent variables used in the analysis are presented in **Error! Reference source not found.**. Out of the 12112 sampled farmers interviewed in the study 23.6% were female and 76.4% were male. The gender distribution is not consistent with Malawi’s population distribution (51% female and 49% Male) (National Statistical Office of Malawi, 2018). This may be a consequence of the differences between farmland ownership between the gender groups. The average age estimated at 46.1 years exhibited bias compared to the national estimates of 17.4 (National Statistical Office of Malawi, 2018). Again, this is a consequence of the recruitment criterion as only adults participated in the study. The average family size of 4.67 fitted approximately the national average of 4.4 (National Statistical Office of Malawi, 2018). On average, a sampled farmer had been to primary school i.e. standard 7. The overall landholding size for the participants was 2.8 acres, which is consistent with the national estimated average landholding size of 2.96 acres ([FAO], 2011). Significant differences were found between landholding sizes between male and female landowners (p=0.007), with males having higher land sizes of 2.9 acres and females with 2.47acres. Globally, women consistently have lower access to land capital, and for those that have access, the landholding size is mostly significantly lower than their male counterparts ([FAO], 2011). |About 88% of the household used own farmland but out these only 1% had legal documents to support their ownership claim. The remaining 11% used rented (7%), borrowed for free (3%), encroached land (<1%), and farm tenant (<1%).

### 2.5 Determinants of adoption of adaptation strategies

Table 1 shows the results from multinomial logit regression used to predict the effect of land tenure arrangements on the adoption of climate change adaptation technologies where the baseline category is no adoption. The model is a good fit for the data used: the Wald test indicates that *x*^2^ = 495.2 and *P* > *x*^2^ = 0.00. This entails that the null hypothesis that all regression coefficients are jointly equal to zero is rejected.

**Table 1.** Descriptive statistics of variables used in the analysis

|  | Mean | SD | Description | Hypothesized impact |
| --- | --- | --- | --- | --- |
| *socio-economic characteristics* |  |  |  |  |
| Age | 46.11 | 15.65 | Age of farmer | (±) |
| sex | 0.76 | 0.42 | Binary, 1 = Male, 0 otherwise | (+) |
| Education | 6.92 | 3.61 | Education level of farmer | (+) |
| *Land parcel characteristics* |  |  |  |  |
| Plot size | 2.79 | 22.69 | Size of plot in acres | (+) |
| Good Soil quality | 0.53 | 0.5 | Binary, 1 = Yes, 0 = Otherwise | (-) |
| Erosion rate low | 0.83 | 0.99 | Binary, 1 = Yes, 0 = Otherwise | (+) |
| *Tenure variables* |  |  |  |  |
| Full ownership | 0.88 | 0.32 | Binary, 1 = Yes, 0 = Otherwise | (+) |
| Rent_Short_Term | 0.07 | 0.26 | Binary, 1 = Yes, 0 = Otherwise | (-) |
| Borrowed_Free | 0.03 | 0.18 | Binary, 1 = Yes, 0 = Otherwise | (-) |
| Encroachment | 0.007 | 0.09 | Binary, 1 = Yes, 0 = Otherwise | (-) |
| Tenant | 0.002 | 0.05 | Binary, 1 = Yes, 0 = Otherwise | (-) |
| *Institutional variables* |  |  |  |  |
| Extension access | 0.48 | 0.5 | Binary, 1 = Yes, 0 = Otherwise | (+) |
| Credit access | 0.29 | 0.45 | Binary, 1 = Yes, 0 = Otherwise | (+) |

**Table 2.** multinomial regression model results for determinants adoption of climate change adaptation strategies

| Variable | ER | SF | SFER |
| --- | --- | --- | --- |
| age | -0.0119*** | -0.00639 | -0.00847** |
|  | -0.0044 | -0.00416 | -0.00401 |
| sex | 0.189 | 0.14 | 0.292** |
|  | -0.155 | -0.147 | -0.141 |
| highestlevel | 0.00191 | -0.00846 | 0.000246 |
|  | -0.0187 | -0.0179 | -0.0172 |
| acres | -0.00618** | -0.00395** | -0.00352*** |
|  | -0.00242 | -0.00156 | -0.00137 |
| soilquality | -0.186* | -0.0992 | -0.200** |
|  | -0.0976 | -0.0925 | -0.0892 |
| erosionrate | -0.237*** | 0.0761 | -0.015 |
|  | -0.0781 | -0.0717 | -0.0695 |
| Rent_Short_Term | -0.0954 | -1.342*** | -1.516*** |
|  | -0.174 | -0.179 | -0.165 |
| Borrowed_Free | 0.391 | -0.715** | -0.824*** |
|  | -0.302 | -0.308 | -0.29 |
| Encroachment | 14.04 | 14.46 | 14.27 |
|  | -905.1 | -905.1 | -905.1 |
| Tenant | 13.58 | 13.62 | 14.83 |
|  | -1,798 | -1,798 | -1,798 |
| access | -0.00985 | 0.146 | 0.477*** |
|  | -0.138 | -0.132 | -0.127 |
| credit_access | -0.0727 | -0.00707 | -0.103 |
|  | -0.145 | -0.138 | -0.133 |
| coupon_access | 0.313* | 0.809*** | 0.692*** |
|  | -0.166 | -0.157 | -0.153 |
| Constant | 2.452*** | 2.290*** | 3.556*** |
|  | -0.352 | -0.334 | -0.323 |

#### 2.5.1 Land tenure

Land tenure variables had a significant effect on the adoption of soil fertility technologies and a combination of fertility and erratic rainfall adaptation strategies. As envisaged Use of rented or borrowed land reduced the probability of adoption technologies. *Rent* and *Borrowed for free* carry negative coefficients in the SF and SFER models implying that using rented land is associated with a declining probability of adoption. The predictive margins reported in Table 3 show that users of land borrowed for free are more likely not to adopt technologies that those using own land i.e. 4.3% as opposed to 2.6%. Likewise, those using rented land are three times more likely (7.2%) not to adopt any technology than those using own land (2.3%). The observation in this study is consistent with findings in South Africa (Gbetibouo, Hassan, & Ringler, 2010). Land tenure arrangements other than ownership are associated with declining security. Thus, the long-term dimension on return in investing in SF and SRER discourages the adoption of such practices among farmers who are using rented land as they may not control the land for long to reap the benefits of their investment (Kpadonou, et al., 2017).

**Table 3.**
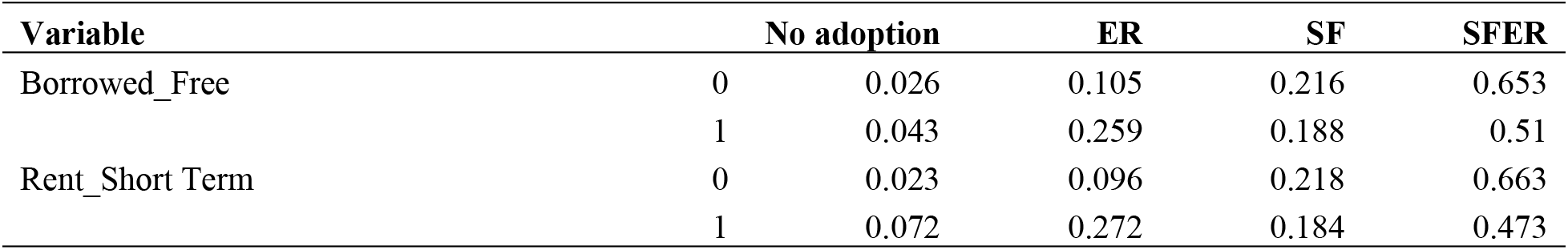
Predictive margins for land tenure variables

#### 2.5.2 Institutional variables

Feder and Slade (1984) describe technology adoption as a multistage process that farm households go through from the time they become aware of innovation to the time that they decide to start using the technology. Central to the adoption decisions is the role of information (Simtowe, Asfaw, & Abate, 2016) and the extension systems are in existence to save this purpose. In line with expectation, access to extension increased the probability of adopting SFER. Interestingly, it did not significantly influence adoption SF and ER. Technology promotion is often done in the realm of soil and water conservation which encompasses both SF and ER technologies as such farmers with access to an effective extension service are likely to adopt a combination of the two groups. Despite 29% of participants reporting access to credit no significant relationship was observed with the adoption of adaptation strategies. A probable explanation could be that not all credit gotten is used for agriculture production (Diagne & Zeller, 2001; Simtowe & Zeller, 2006).

### 2.6 Land parcel characteristics

Plot size was also found to significantly affect adoption. As the size of farmland increases, the likelihood of adoption ER, SF and SFER decreases. Several studies found that farmers who have big farms are more like to invest in SWC (Habtamu, 2006; Mohammed et al., 2018) highlighting that most farmers with large land sizes are older an often lack labor required for maintaining conservation structures. On the contrary, Lovo (2016) that increasing land size increases the probability of adoption. Almost half (47%) of study participants perceived their soils to be of poor quality. Naturally, if acting rationally these households are expected to invest in SF technologies. Interestingly, The dummy variable for soil quality had a positive and significant coefficient in the ER and ERSF and insignificant for SF. This result suggests that farmers that adopted such technologies were not responding to their perception of soil status. This result is worrisome as it raises the potential for land degradation of fertile lands as farmers are not responsive to conditions.

### 2.7 Socio-economic characteristics

Sex of the farmer exhibited a positive and significant relationship with the adoption of ER implying that a male farmer has a higher chance of adoption than their female counterparts. The observed differences between male and female farmers do not emanate from gender but differences in access to resources (Doss & Morris, 2001). Age carries a negative coefficient indicating that older farmers are less likely to adopt. This finding agrees with (Farid, Tanny, & Suppadit, 2015; Kamau, Smale, & Mutua, 2014) but disagrees with Mango, Makate, Tamene, Mponela, and Ndengu (2017) who found a positive correlation between age and adoption.

## 3 Conclusion

The definitive aim of this study was to bring to fore the evidence of the importance of tenure considerations in the designing, development, and implementation of agricultural investment programs and projects. This was done by analyzing how land tenure and other determinants affect the adoption of climate adaptation technologies in Malawi. Land tenure has shown to significantly affect adoption in varying ways depending on the technology in question. The evidence generated from this study not only confirms that the provision of secure ownership in Malawi can increase the adoption of technologies significantly, but also that failure to provide this security negatively impacts the production and income benefits associated with the adoption of agricultural investments under secure land ownership.

The proliferation of borrowed or rented in Malawi’s agricultural sector, 10% of all cultivated land in 2016 (National Statistical Office of Malawi [NSO], 2017), entails that due consideration should be put to advising landowners to either invest in climate change adaptation themselves or set conditions within the tenancy agreements to avoid land degradation. As the land policy reforms gather pace it is important for extension on land matters to create awareness on the same.

